# Amphitrophic *Listeria monocytogenes* causes one-third of invasive listeriosis yet remains undetected by clonal complex-based risk classification

**DOI:** 10.64898/2026.03.28.715028

**Authors:** Javier Gamboa

## Abstract

Conventional genomic risk classification of *Listeria monocytogenes* assigns clonal complexes to hypervirulent (CC1, CC2, CC4, CC6) or hypovirulent (CC9, CC121) categories based on population-level frequency ratios, leaving all remaining diversity in an undifferentiated “intermediate” category that carries no defined risk assessment. We analysed 436 genomes from confirmed invasive listeriosis across 19 countries using multi-dimensional genomic profiling of virulence and persistence determinants and demonstrate that this approach systematically misclassifies a major fraction of clinically relevant *L. monocytogenes*. Amphitrophic lineages — carrying simultaneous genomic competence for clinical virulence (functional *inlA*, mean virulence score 52.7 +/- 6.6) and industrial persistence (SSI-1 in 94.1%, mean persistence score 66.8 +/- 11.6) — constitute 31.0% of invasive disease, within 3.6 percentage points of the established hypervirulent category (34.6%). Of these 135 amphitrophic clinical isolates, 91.1% were classified as “intermediate” under conventional taxonomy. The five principal amphitrophic CCs (CC8, CC7, CC3, CC5, CC88) appear with indistinguishable dual-fitness genotypes in both clinical and food-chain datasets, establishing that the same organisms persist in processing facilities and cause invasive human disease. Decomposition of the species-level virulence-persistence trade-off (rho = -0.523) by trophic strategy reveals it to be a Simpson’s paradox: no within-strategy correlation is significantly negative, and the only significant signal is a positive amphitrophic correlation (rho = +0.221, p = 0.010) indicating synergy rather than trade-off. Multi-dimensional profiling increases risk-stratified detection from 32.3% (conventional) to 65.6% of clinical isolates — a 103% improvement. These findings demonstrate that clonal complex identity alone leaves one-third of clinically significant *L. monocytogenes* uncharacterised, and that effective One Health genomic surveillance requires simultaneous assessment of virulence and persistence at the isolate level.

## 1. Introduction

Whole genome sequencing has become the definitive tool for *Listeria monocytogenes* surveillance, underpinning outbreak detection, source attribution and regulatory compliance across the food chain and clinical microbiology (Moura et al., 2016). Integrated WGS-based surveillance across food, environmental and clinical sectors is now mandated in multiple jurisdictions, yet the interpretive infrastructure to support cross-sector integration does not exist. Clinical and food safety laboratories analyse identical genomes through incompatible conceptual frameworks: one oriented toward virulence and outbreak linkage, the other toward persistence and source control.

This disconnect is anchored in the virulence–persistence trade-off model, which holds that *L. monocytogenes* lineages specialise toward either clinical pathogenesis or environmental survival (Maury et al., 2016). Under this model, clonal complexes CC1, CC2, CC4 and CC6 — designated hypervirulent based on their overrepresentation in invasive listeriosis relative to food/environmental reservoirs — account for the majority of severe clinical disease. Complexes CC9 and CC121 — designated hypovirulent based on their dominance in food processing environments and rarity in clinical cases — are treated as low clinical risk. All remaining diversity is assigned to an “intermediate” category defined not by biological characterisation but by absence of extreme frequency bias in either direction.

We recently demonstrated that this tripartite classification conceals critical biological structure (Gamboa, 2026). By integrating virulence, persistence, clonality and antimicrobial resistance determinants into a multi-dimensional genomic assessment — the Genomic Intelligence Framework (GIF) — we resolved 903 food-chain genomes into three trophic strategies based on genomic functional capacity rather than population frequency: nosotrophic lineages specialised for acute host invasion, saprotrophic lineages adapted for environmental persistence, and amphitrophic lineages retaining simultaneous competence for both. The amphitrophic category — comprising 39.1% of food-chain isolates and dominated by CC7, CC3, CC8, CC5 and CC88 — directly challenged the trade-off as a universal biological constraint by demonstrating coexistence of functional *inlA*, stress survival islands and persistence determinants within individual genomes.

However, the initial characterisation was confined to food-chain isolates. The clinical relevance of amphitrophic lineages — whether they actually cause human disease, and if so at what frequency — remained an open question with direct consequences for risk assessment and surveillance policy. If amphitrophic organisms are genuinely dual-fitness, they should appear in invasive listeriosis at substantial frequency. If the trade-off model holds at the biological level, they should be rare or absent in clinical presentation.

Here we test whether amphitrophic lineages cause invasive human disease, applying multi-dimensional genomic profiling to 436 genomes from confirmed invasive listeriosis across 19 countries. The results expose a systematic blind spot in current risk assessment: amphitrophic lineages constitute nearly one-third of invasive disease, virtually matching the frequency of established hypervirulent lineages, yet were invisible to conventional classification.

## 2. Materials and Methods

### 2.1 Clinical dataset assembly

*L. monocytogenes* isolates from confirmed invasive listeriosis were retrieved from NCBI Pathogen Detection (accessed March 2026). Inclusion criteria required isolation type “clinical”, GenBank assembly accession (GCA_*), assigned SNP cluster and confirmed human source. From 20,219 qualifying records, 438 isolates were selected through stratified sampling to ensure proportional representation across serogroups 4b (enriched for CC1/CC2/CC4/CC6), 1/2a (CC7/CC8/CC155/CC121), 1/2b (CC3/CC5), 1/2c (CC9) and unknown serovar with confirmed invasive source (blood, cerebrospinal fluid, placenta). Geographic representation was enforced through per-country caps of 30% and a minimum of 10 countries. Two assemblies failed download, yielding 436 genomes for analysis.

Metadata were extracted from NCBI Pathogen Detection: collection date (available for 98.9% of isolates; range 1981–2025), isolation source, location, SNP cluster, virulence genotypes, stress genotypes and AMR genotypes. Isolation sources comprised blood (30.5%), unspecified clinical/human (65.7%), cerebrospinal fluid (3.0%) and placenta (0.7%). All isolates were independently confirmed as clinical by NCBI Pathogen Detection curation pipeline.

### 2.2 GIF processing and trophic classification

All 436 assemblies were processed through the GIF v1.0 pipeline as previously described (Gamboa, 2026): cgMLST allele calling against the Institut Pasteur 1,748-locus scheme via chewBBACA, virulence and resistance gene detection via ABRicate, MLST/CC assignment, and hierarchical clustering against the NCBI Pathogen Detection reference database. Component scores (V, P, C, R) were computed under both industrial (V:30/P:40/C:20/R:10) and clinical (V:40/P:20/C:30/R:10) calibrations. Trophic strategy was assigned per isolate following established thresholds (Gamboa, 2026): nosotrophic (high V-Score, low P-Score), saprotrophic (low V-Score, high P-Score), amphitrophic (both scores above respective thresholds), unassigned (neither criterion met).

P-Score Level 2 (operational persistence based on temporal facility data) was estimated via Bayesian imputation from genetic persistence markers for all clinical isolates, as temporal metadata are unavailable for clinical cases. This approach yields conservative estimates validated against phenotypic persistence classification (AUC = 0.980 in the Fagerlund longitudinal dataset; Gamboa, 2026).

### 2.3 Comparison with conventional classification

Each isolate was assigned a Maury epidemiological category based on CC identity: hypervirulent (CC1, CC2, CC4, CC6), hypovirulent (CC9, CC121) or intermediate (all remaining CCs), following Maury et al. (2016). Cross-tabulation against GIF trophic assignment quantified concordance and divergence. The proportion of clinical isolates flagged as “high risk” was compared between conventional classification (hypervirulent CCs only) and GIF (nosotrophic + amphitrophic).

### 2.4 Cross-context comparison

Clinical trophic distributions, V–P correlations, calibration differentials and CC-level scores were compared with the four food-chain datasets from Gamboa (2026): Fagerlund (n = 513, Norwegian food processing), Stasiewicz (n = 191, US retail), Zhang (n = 151, Beijing clinical) and Kurpas (n = 48, Polish RTE meat). NCBI-derived virulence genotypes and stress genotypes provided independent validation of GIF component assignments.

### 2.5 Statistical analysis

Spearman rank correlations were computed for V-Score versus P-Score globally and stratified by trophic strategy. Descriptive statistics were calculated for all GIF components by trophic category, CC and Maury class. All analyses were performed in Python 3.12 with complete reproducibility code deposited at https://github.com/jgamboa-biotecno/GIF-Framework.

## 3. Results

### 3.1 Dataset overview

Multi-dimensional genomic profiling processed all 436 clinical assemblies with 100% completion, identifying 57 distinct clonal complexes distributed across Lineage I (56.2%) and Lineage II (41.5%). The dataset encompasses 19 countries (UK 18.0%, Germany 18.0%, USA 16.4%, Netherlands 14.1%, 15 additional countries) with collection dates spanning 1981–2025.

### 3.2 The amphitrophic blind spot: one-third of invasive listeriosis invisible to conventional classification

GIF classified clinical isolates into three trophic strategies whose frequencies diverge sharply from population-level expectations (Table 1, Fig. 1).

**Table 1.** Trophic strategy distribution in clinical versus food-chain context.

| Strategy | n<br>(clinical) | %<br>clinical | % food-chain | Enrichment | V-Score | P-Score | InlA<br>functional | SSI-1 |
| --- | --- | --- | --- | --- | --- | --- | --- | --- |
| Nosotrophic | 151 | 34.6 | 22.7 | 1.53× | 76.7 ±<br>8.5 | 30.2 ±<br>1.0 | 100% | 0% |
| Amphitrophic | 135 | 31.0 | 39.1 | 0.79× | 52.7 ±<br>6.6 | 66.8 ±<br>11.6 | 100% | 94.1% |
| Saprotrophic | 51 | 11.7 | 5.8 | 2.02× | 13.7 ±<br>8.4 | 72.3 ±<br>13.4 | 0% | 90.2% |
| Unassigned | 99 | 22.7 | 32.4 | 0.70× | 50.0 ±<br>5.6 | 30.6 ±<br>1.5 | 100% | 0% |

The central finding is the near-parity between nosotrophic and amphitrophic clinical frequencies: 34.6% versus 31.0%, a difference of 3.6 percentage points. Nosotrophic dominance of clinical presentation is expected and well-characterised — these are the established hypervirulent lineages (CC1, CC2, CC4, CC6, CC87, CC14) that the field has recognised since Maury et al. (2016). The amphitrophic frequency, however, represents a previously unquantified disease burden: 135 cases of confirmed invasive listeriosis caused by organisms whose defining characteristic — simultaneous virulence and persistence competence — placed them outside all existing risk categories.

Under conventional classification, these 135 amphitrophic clinical isolates were categorized as: “intermediate” (123, 91.1%), “hypovirulent” (6, 4.4%), unresolved CC (5, 3.7%), and — notably — “hypervirulent” (1, 0.7%). The single hypervirulent amphitroph is a CC2 isolate (V-Score 63.3, P-Score 68.3) carrying functional *inlA* and high persistence despite belonging to a lineage conventionally assumed to lack environmental fitness. The “intermediate” designation carries no defined clinical action and no specified industrial significance — it is a residual category indicating that a lineage’s frequency ratio does not reach the threshold for either hypervirulent or hypovirulent designation. In practice, 95.6% of amphitrophic clinical isolates were classified into categories that communicate nothing about their dual-fitness capability (91.1% “intermediate” + 4.4% “hypovirulent”), while the remaining 3.7% could not be classified at all due to unresolved CC assignment.

### 3.3 Quantifying the detection gap

Conventional risk classification (Maury hypervirulent CCs only) flags 141 of 436 clinical isolates (32.3%) as high risk. GIF, by resolving the “intermediate” category into biologically defined populations, identifies 286 isolates (65.6%) as carrying significant risk — 151 nosotrophic (acute clinical threat) plus 135 amphitrophic (dual clinical–industrial threat). This represents a 103% increase in risk-stratified detection (Table 2, Fig. 3).

**Table 2.** Risk detection: conventional classification versus GIF.

| System | High-risk isolates | % of clinical dataset | Basis |
| --- | --- | --- | --- |
| Conventional (Maury) | 141 | 32.3% | CC1, CC2, CC4, CC6 frequency ratio |
| GIF — nosotrophic | 151 | 34.6% | Genomic virulence profile |
| GIF — amphitrophic | 135 | 31.0% | Genomic dual-fitness profile |
| **GIF — combined** | **286** | **65.6%** | Virulence and/or persistence competence |
| **Additional detection** | **+145** | **+103%** | Amphitrophs + reclassified nosotrophs |

The additional detection by GIF comprises two populations not flagged by conventional classification. First, amphitrophic isolates invisible to conventional classification because their CCs (CC8, CC7, CC3, CC5, CC155, CC88, CC224) fall into the undifferentiated “intermediate” bin. Second, nosotrophic isolates — CC87 (n = 12, V-Score 83.3) and CC14 (n = 7, V-Score 70.0) — whose CCs lack sufficient representation in European surveillance to reach Maury’s hypervirulent frequency threshold, despite carrying genomic virulence profiles indistinguishable from CC1/CC4.

### 3.4 Amphitrophic dual-fitness genotype preserved across clinical and industrial contexts

The five principal amphitrophic CCs identified in food-chain environments (CC8, CC7, CC3, CC5, CC88; Gamboa, 2026) are all recovered in clinical isolates with identical dual-fitness genotypes: 100% functional *inlA*, 94.1% SSI-1, 10.4% *bcrABC*, 5.2% *qacH* (Table 3). Additionally, CC155 (n = 12) and CC224 (n = 8) emerge as amphitrophic lineages in clinical context, expanding the confirmed amphitrophic repertoire.

**Table 3.** Amphitrophic CCs: clinical versus food-chain genotype.

| CC | n (clinical) | V-Score | P-Score | *inlA* | SSI-1 | *bcrABC* | Confirmed in food-chain |
| --- | --- | --- | --- | --- | --- | --- | --- |
| CC8 | 30 | 48.7 | 64.2 | 100% | 100% | 0% | Yes (Fagerlund) |
| CC7 | 18 | 58.2 | 66.7 | 100% | 100% | 6% | Yes (Fagerlund) |
| CC155 | 12 | 48.1 | 73.4 | 100% | 100% | 25% | **New** |
| CC3 | 12 | 68.0 | 75.1 | 100% | 100% | 33% | Yes (Stasiewicz) |
| CC5 | 11 | 56.7 | 71.5 | 100% | 100% | 18% | Yes (Stasiewicz) |
| CC88 | 4 | 46.7 | 71.4 | 100% | 100% | 25% | Yes (Fagerlund) |
| CC224 | 8 | 54.2 | 61.9 | 100% | 100% | 0% | **New** |

The genotypic identity between clinical and food-chain amphitrophs is the critical observation. These are not organisms that acquire virulence upon entering a clinical context or persistence upon entering a food-processing context. Their dual capacity is constitutive, encoded in a stable genomic architecture that functions across ecological boundaries. A CC8 isolate recovered from blood culture and a CC8 isolate recovered from a salmon processing environment carry the same *inlA* allele and the same SSI-1 island.

### 3.5 Virulence–persistence trade-off: an ecological artefact resolved by trophic classification

The aggregate V–P correlation across all 436 clinical isolates is rho = −0.523 (p < 0.001). Decomposition by trophic strategy reveals that this apparent trade-off is driven entirely by between-group structure (Table 4, Fig. 2).

**Table 4.** V–P correlation decomposed by trophic strategy.

| Stratum | n | $\rho$ (V, P) | p-value | Interpretation |
| --- | --- | --- | --- | --- |
| All clinical isolates | 436 | −0.523 | <0.001 | Apparent trade-off |
| Nosotrophic only | 151 | −0.085 | 0.297 | Non-significant |
| **Amphitrophic only** | **135** | **+0.221** | **0.010** | **Significantly positive (synergistic)** |
| Saprotrophic only | 51 | −0.035 | 0.807 | Non-significant |

Within trophic strategies, V–P correlations are either non-significant (nosotrophic rho = −0.085, p = 0.30; saprotrophic rho = −0.035, p = 0.81) or significantly positive (amphitrophic rho = +0.221, p = 0.010). The only stratum exhibiting a statistically significant V–P correlation is amphitrophic — and it is positive, indicating that virulence and persistence determinants co-segregate rather than oppose each other. The species-level negative correlation is a Simpson’s paradox artefact produced by mixing three populations with different mean V and P values. Trophic classification resolves this artefact by assigning ecological strategy at the individual isolate level.

This finding, combined with the food-chain data (Gamboa, 2026), provides the fifth independent replication of the V–P negative correlation at species level across datasets spanning multiple countries, three decades and five ecological contexts (Table 5) — and simultaneously demonstrates that this correlation does not reflect a constraint operating on individual genomes.

**Table 5.** V–P correlation across all GIF-characterised datasets.

| Dataset | Context | n | rho (V, P) |
| --- | --- | --- | --- |
| Fagerlund | Food processing (Norway) | 513 | −0.205 |
| Stasiewicz | Retail (USA) | 191 | −0.524 |
| Zhang | Clinical (China) | 151 | −0.618 |
| Kurpas | RTE meat (Poland) | 48 | −0.713 |
| **Clinical** | **Invasive listeriosis (19 countries)** | **436** | **−0.523** |
| **Combined** | **One Health** | **1,339** | **—** |

### 3.6 Saprotrophic clinical enrichment: host vulnerability quantified

Saprotrophic lineages (CC9, CC121) appear in invasive listeriosis at 2.02-fold their food-chain frequency (11.7% vs 5.8%). GIF assigns these isolates appropriately low V-Scores (13.7 ± 8.4): 87% of CC9 clinical isolates carry truncated *inlA*, none harbour LIPI-3, and their intrinsic invasion potential is minimal. Simultaneously, P-Scores are high (72.3 ± 13.4): 100% of clinical CC9 carry SSI-1, 15.7% carry *qacH*, and 15.7% carry *bcrABC*.

This dual assessment — low intrinsic virulence, high persistence potential — provides the mechanistic interpretation that CC-level classification cannot: these organisms reach clinical presentation through host-side vulnerability (immunocompromise, extremes of age, pregnancy) or dose-dependent exposure via persistent environmental contamination, not through intrinsic pathogenic capability. The GIF clinical score (27.7 ± 4.5, LOW tier) correctly communicates this distinction.

### 3.7 Maury “intermediate” decomposed: from residual category to defined biology

Cross-tabulation of all 234 Maury “intermediate” clinical isolates against GIF trophic classification reveals that this epidemiological category masks four genomically distinct populations (Table 6, Fig. 4).

**Table 6.**
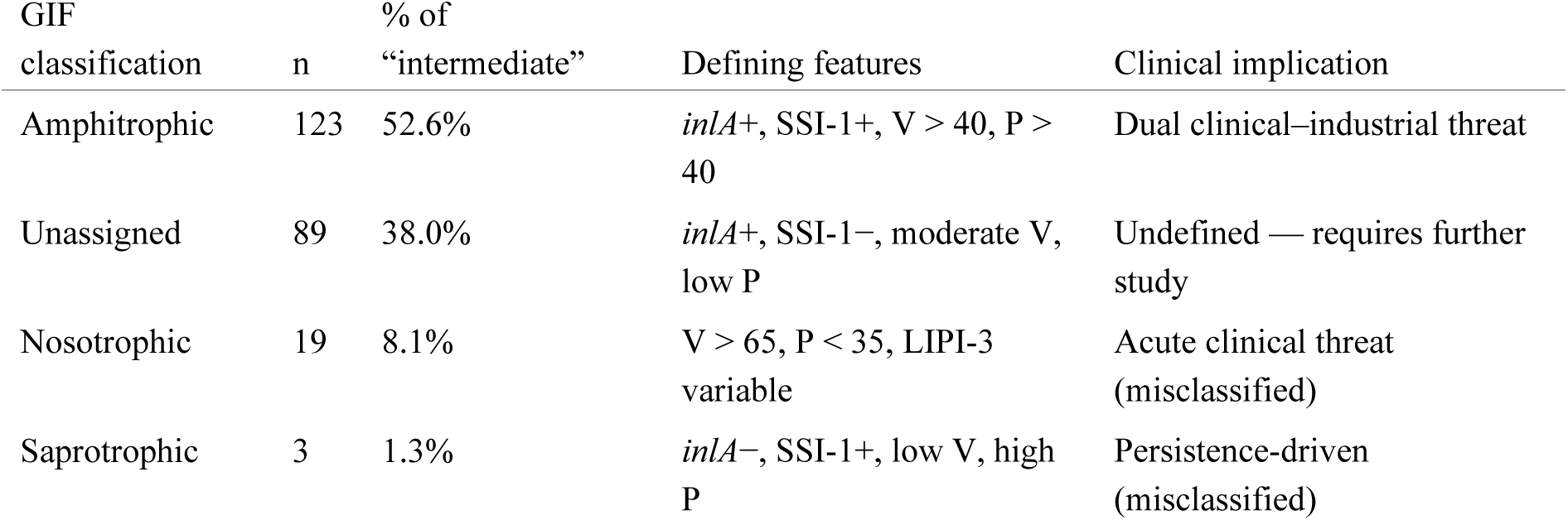
Genomic decomposition of Maury “intermediate” category.

| GIF classification | n | % of “intermediate” | Defining features | Clinical implication |
| --- | --- | --- | --- | --- |
| Amphitrophic | 123 | 52.6% | *inlA*+, SSI-1+, V > 40, P > 40 | Dual clinical–industrial threat |
| Unassigned | 89 | 38.0% | *inlA*+, SSI-1–, moderate V, low P | Undefined — requires further study |
| Nosotrophic | 19 | 8.1% | V > 65, P < 35, LIPI-3 variable | Acute clinical threat (misclassified) |
| Saprotrophic | 3 | 1.3% | *inlA*–, SSI-1+, low V, high P | Persistence-driven (misclassified) |

The distinction between Maury’s “intermediate” and GIF’s “amphitrophic” is not terminological — it is conceptual. Maury’s classification answers “where does this lineage predominantly appear?” by computing clinical-to-food frequency ratios across surveillance databases. GIF’s classification answers “what can this genome do?” by interrogating functional markers in the individual isolate. The former requires population-level data and cannot classify novel or underrepresented lineages; the latter operates on a single genome regardless of surveillance representation. Nearly half (47.4%) of Maury’s “intermediate” isolates are not amphitrophic: 19 are genomic nosotrophs (CC87, CC14) whose CCs lack European surveillance representation to cross Maury’s frequency threshold despite carrying virulence profiles functionally equivalent to CC1/CC4, and 89 are lineages without defined persistence markers that require further characterisation.

### 3.8 Dual calibration empirically validated

The clinical calibration (V:40/P:20/C:30/R:10) shifts GIF scores relative to the industrial calibration (V:30/P:40/C:20/R:10) in the biologically expected direction for each trophic strategy (Table 7).

**Table 7.** Calibration differential by trophic strategy.

| Strategy | GIF Industrial | GIF Clinical | Delta | Expected direction |
| --- | --- | --- | --- | --- |
| Nosotrophic | 40.5 | 44.7 | **+4.1** | Elevated (primary clinical threat) |
| Amphitrophic | 48.4 | 43.0 | −5.4 | Reduced (persistence less weighted) |
| Saprotrophic | 38.4 | 27.7 | **−10.7** | Strongly reduced (low intrinsic virulence) |

The clinical calibration elevates nosotrophic risk (acute invasion specialists) by 4.1 points and depresses saprotrophic risk (low intrinsic virulence) by 10.7 points. Amphitrophic isolates score higher under industrial calibration (+5.4 points) where persistence receives greater weight, reflecting their dual relevance to food safety. This context-dependent score behaviour — identical genome, different risk interpretation depending on setting — is precisely the capability required for One Health surveillance operating across the clinical–industrial boundary.

## 4. Discussion

### 4.1 The amphitrophic imperative for One Health surveillance

The defining result of this study is that 135 confirmed invasive listeriosis cases — 31.0% of the clinical dataset, within 3.6 percentage points of the established hypervirulent frequency — were caused by amphitrophic lineages that conventional classification either ignored or actively misclassified. These are not marginal organisms or statistical curiosities. They constitute a disease burden comparable in magnitude to the hypervirulent lineages that have anchored *L. monocytogenes* risk assessment for a decade, yet they were systematically invisible because the dominant classification framework lacks the biological resolution to detect them.

The implications extend beyond clinical microbiology. The same amphitrophic CCs (CC8, CC7, CC3, CC5, CC88) that appear in 31% of invasive listeriosis also dominate food processing environments (39.1% of food-chain isolates). They carry identical dual-fitness genotypes in both contexts: functional *inlA* enabling intestinal invasion, SSI-1 enabling environmental stress survival, and in a subset *bcrABC* enabling biocide tolerance. A CC8 isolate from a blood culture in Germany and a CC8 isolate from a salmon processing facility in Norway are not analogous organisms from parallel ecological niches — they are the same biological population operating across the clinical–environmental boundary.

Under current practice, these two isolates are managed by different agencies, classified under different frameworks, and stored in different databases. The clinical CC8 is a listeriosis case; the industrial CC8 is a food safety non-conformity. Neither system recognises their shared identity or the chain connecting them. Multi-dimensional trophic classification, by assigning identical ecological strategy to both and flagging the dual-fitness genotype, provides the interpretive bridge that One Health genomic surveillance requires.

Consider the practical difference. A CC8 blood isolate classified as “intermediate” under Maury triggers standard clinical management and routine notification. No industrial investigation is initiated because “intermediate” carries no signal of environmental relevance. The processing facility harbouring this clone continues to operate and contaminate. Under trophic classification, the same isolate is identified as amphitrophic with explicit persistence markers (SSI-1+, P-Score 64.2). This triggers cross-referencing against NCBI Pathogen Detection — where 82 unique SNP clusters from amphitrophic clinical isolates in this dataset alone represent potential bridges to food-chain sources. If a match is identified, the facility is targeted for intervention before further clinical cases occur.

The reverse scenario is equally consequential. A CC7 environmental positive classified as “intermediate” in a food processing facility triggers routine cleaning and recontrols — the standard response for a lineage with no defined risk profile. Under trophic classification, CC7 is amphitrophic (V-Score 58.2, P-Score 66.7, functional *inlA*, SSI-1+). This triggers aggressive eradication rather than routine management, and cross-referencing with clinical surveillance to determine whether human cases attributable to this clone have already occurred.

### 4.2 Resolving the trade-off artefact: from population statistics to individual genomics

The significantly positive intra-amphitrophic V–P correlation (rho = +0.221, p = 0.010) in clinical isolates, combined with the absence of significant trade-off within either nosotrophic (rho = −0.085, p = 0.30) or saprotrophic (rho = −0.035, p = 0.81) strata, provides the most direct evidence to date that the virulence–persistence trade-off does not constrain individual *L. monocytogenes* genomes. Combined with food-chain data (Gamboa, 2026), this observation has now been replicated across six datasets spanning clinical, food processing, retail and environmental contexts.

The species-level negative V–P correlation (rho = −0.523 in clinical isolates, −0.205 to −0.713 across food-chain datasets) reflects the coexistence of ecologically distinct populations — nosotrophic, amphitrophic and saprotrophic — whose different mean V and P values produce a Simpson’s paradox when pooled. The trade-off vanishes entirely within strategies: no stratum shows a significant negative V–P correlation. The only significant within-strategy signal is the amphitrophic positive correlation — synergy, not trade-off.

This distinction has immediate practical consequences. The trade-off model predicts that persistent industrial strains are inherently low clinical risk and that virulent clinical strains are self-limiting in processing environments. The amphitrophic model predicts that neither assumption holds for approximately one-third of the *L. monocytogenes* population. Risk assessment systems calibrated on the trade-off assumption will systematically underestimate the threat from amphitrophic lineages in both clinical and industrial settings — as our data demonstrate they currently do.

### 4.3 Saprotrophic clinical isolates: what V-Score captures that CC cannot

The 2.02-fold enrichment of saprotrophic lineages in clinical context (11.7% vs 5.8% food-chain) demands mechanistic interpretation. The assignment of low V-Score (13.7) to CC9 clinical isolates — despite their clinical origin — captures the biological reality that these organisms reached invasive disease not through intrinsic virulence but through host vulnerability or dose-dependent exposure via persistent contamination. The simultaneously high P-Score (72.3) identifies the environmental pathway: these are organisms whose persistence in food processing (100% SSI-1, 15.7% *qacH*) creates the exposure conditions under which even a hypovirulent strain can cause disease in susceptible hosts.

This dual assessment is impossible under conventional classification, which assigns a single risk label (“hypovirulent”) that correctly describes virulence potential but obscures the persistence-driven exposure pathway. A binary system must either flag CC9 as high risk (overestimating intrinsic virulence) or dismiss it as low risk (ignoring the persistence mechanism that enables clinical presentation). Multi-dimensional scoring resolves this tension by communicating both dimensions simultaneously.

### 4.4 From epidemiological frequency to genomic function

The decomposition of Maury’s “intermediate” category reveals that epidemiological frequency ratios and genomic functional classification capture fundamentally different aspects of *L. monocytogenes* biology. Only 52.6% of “intermediate” isolates are amphitrophic — organisms with demonstrable dual-fitness capacity. The remainder comprises genomic nosotrophs misclassified because their CCs (CC87, V-Score 83.3; CC14, V-Score 70.0) lack sufficient representation in the surveillance databases from which frequency ratios are derived, and functionally unresolved lineages (38.0%) without clear persistence markers that require further characterisation.

This divergence is not a limitation of Maury’s framework within its intended scope — population-level epidemiology — but a demonstration that population-level frequency ratios are insufficient for isolate-level risk assessment. A CC87 isolate causing meningitis carries a virulence profile functionally equivalent to CC4 (V-Score 83.3 vs 94.3), but because CC87’s clinical-to-food ratio has not been established with sufficient statistical power in European surveillance, it is classified as “intermediate” rather than hypervirulent. Trophic classification identifies it as nosotrophic based on its genome, regardless of surveillance representation, because the question relevant to the clinician managing this patient is “what can this organism do?” not “how often does this CC appear in French surveillance databases?”

### 4.5 Limitations

P-Score Level 2 was estimated via Bayesian imputation for all clinical isolates due to absence of facility temporal metadata, providing conservative persistence estimates. Clinical outcome data (severity, mortality, host immunocompromise) were unavailable, precluding individual-level outcome prediction. The dataset is geographically weighted toward Europe and North America (66.5% from Germany, UK, USA and Netherlands). Collection dates span 1981–2025 but concentrate in 2010–2020. The amphitrophic frequency of 31.0% represents a lower bound estimate, as it does not include the 22.7% of unassigned isolates (moderate V-Score, low P-Score) that may harbour persistence capacity undetectable by current genetic markers.

## 5. Conclusions

Applied to 436 confirmed invasive listeriosis genomes, multi-dimensional genomic profiling reveals that amphitrophic *L. monocytogenes* — lineages carrying simultaneous genomic competence for clinical virulence and industrial persistence — constitutes 31.0% of invasive disease, within 3.6 percentage points of the hypervirulent category that has defined *L. monocytogenes* risk assessment since 2016. Of these 135 amphitrophic clinical isolates, 91.1% were classified as “intermediate” under conventional taxonomy — a residual designation that communicates no clinical risk and triggers no industrial investigation. Simultaneous assessment of virulence and persistence increases risk-stratified detection from 32.3% to 65.6% of clinical isolates, a 103% improvement, by resolving this residual category into genomically defined populations with distinct risk profiles and management requirements.

The identification of amphitrophic lineages operating across the clinical–industrial boundary — carrying indistinguishable dual-fitness genotypes in both blood cultures and food processing environments — establishes the biological rationale for integrated One Health genomic surveillance and defines its core requirement: a single interpretive framework capable of simultaneously assessing virulence and persistence at the isolate level. With 1,339 genomes now characterised across five ecological contexts, the multi-dimensional approach validated here provides such a framework. The amphitrophic blind spot demonstrated here — one-third of invasive listeriosis caused by organisms invisible to conventional risk classification — is not a gap in surveillance data but a gap in interpretive capacity. Closing it requires moving beyond population-frequency epidemiology to genome-level functional assessment.

## Ethics Declaration

This study analysed publicly available genome assemblies deposited in NCBI Pathogen Detection by original submitting laboratories. No human subjects were recruited, no patient-identifiable information was accessed, and no clinical samples were collected or processed. All metadata were obtained from public databases. Ethical approval was not required for this retrospective analysis of publicly available genomic data.

## Supporting information

Complete results

GIF vs CCs Statistics

Maury's vs GIF cross

## Data Availability

Complete GIF results for all 436 clinical isolates (152 variables per isolate) are deposited as Supplementary Table S1. Source assemblies and metadata are publicly available from NCBI Pathogen Detection under individual accession numbers listed in Supplementary Table S2. GIF Framework v1.0 specification, scoring algorithms and reference databases: https://github.com/jgamboa-biotecno/GIF-Framework (CC BY-NC 4.0).

---

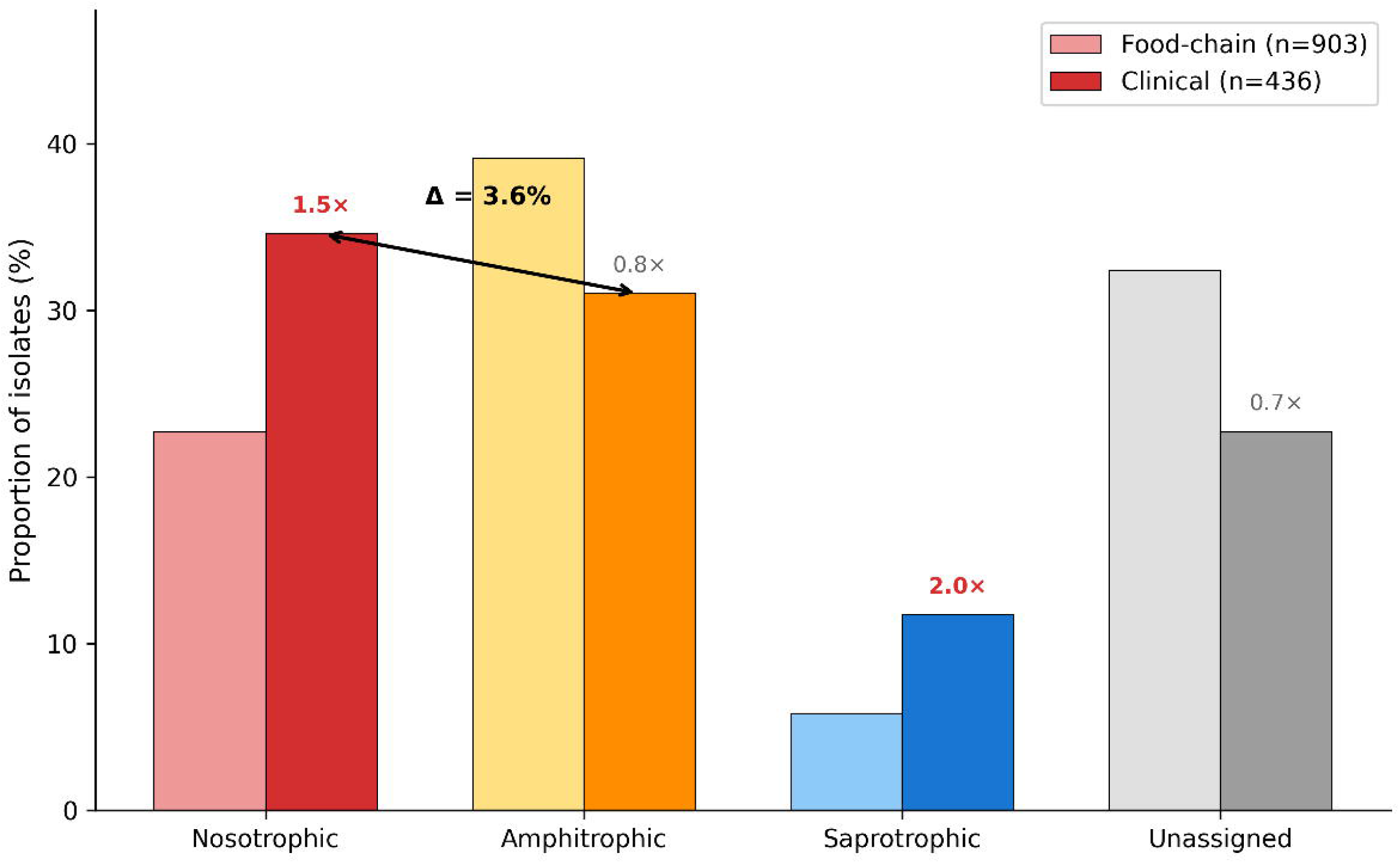

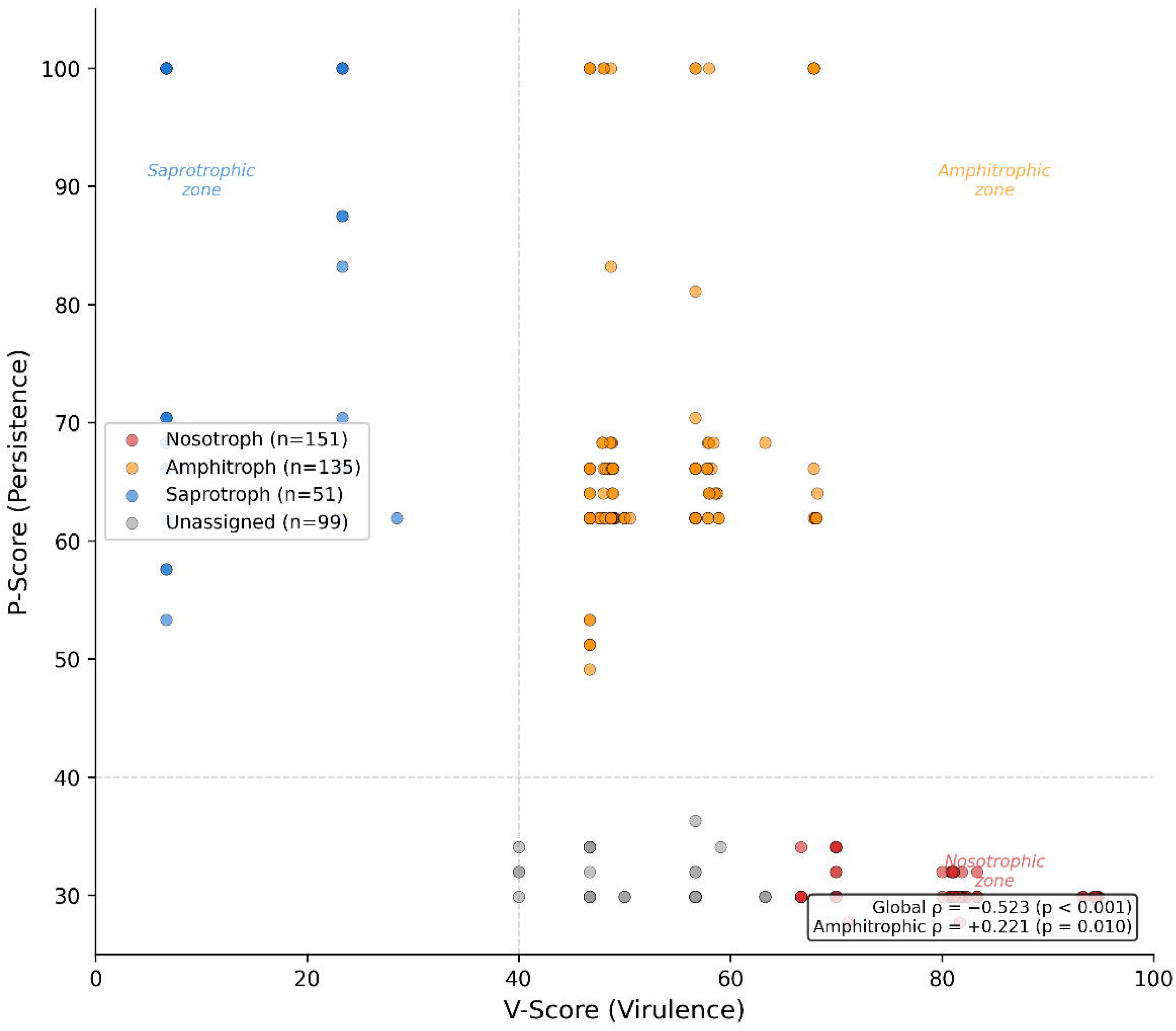

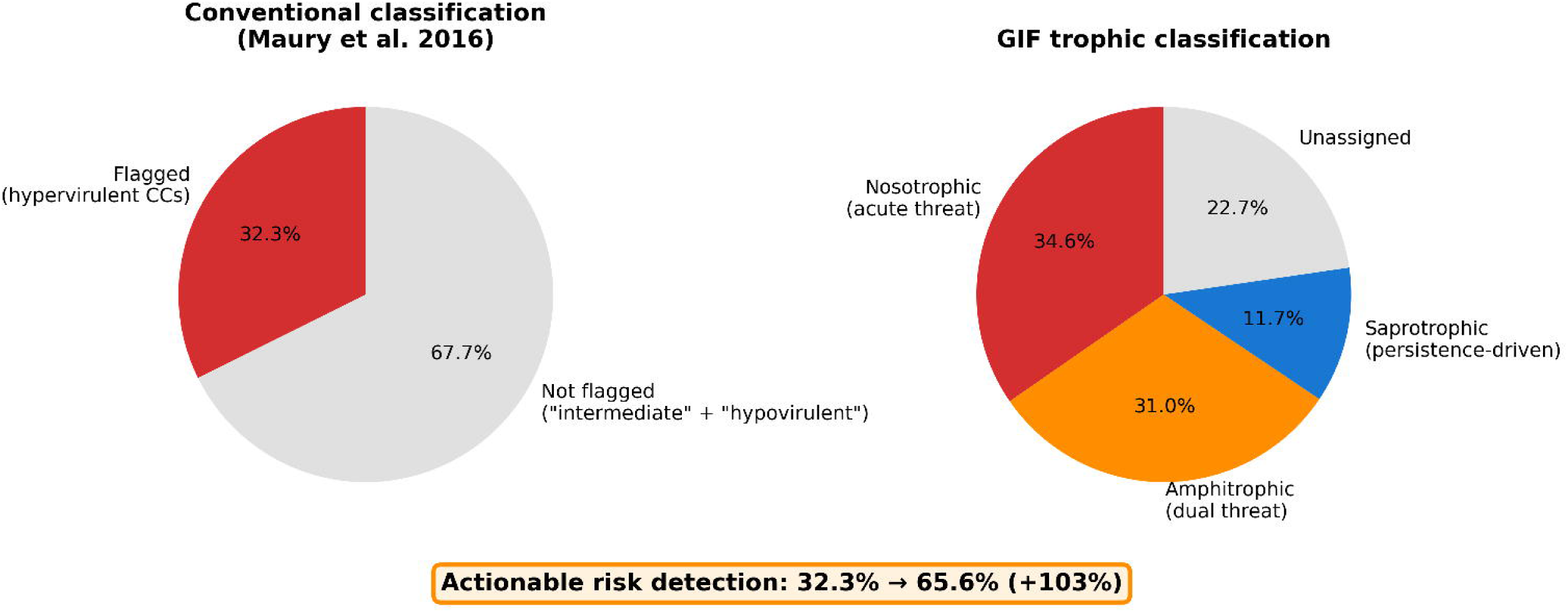

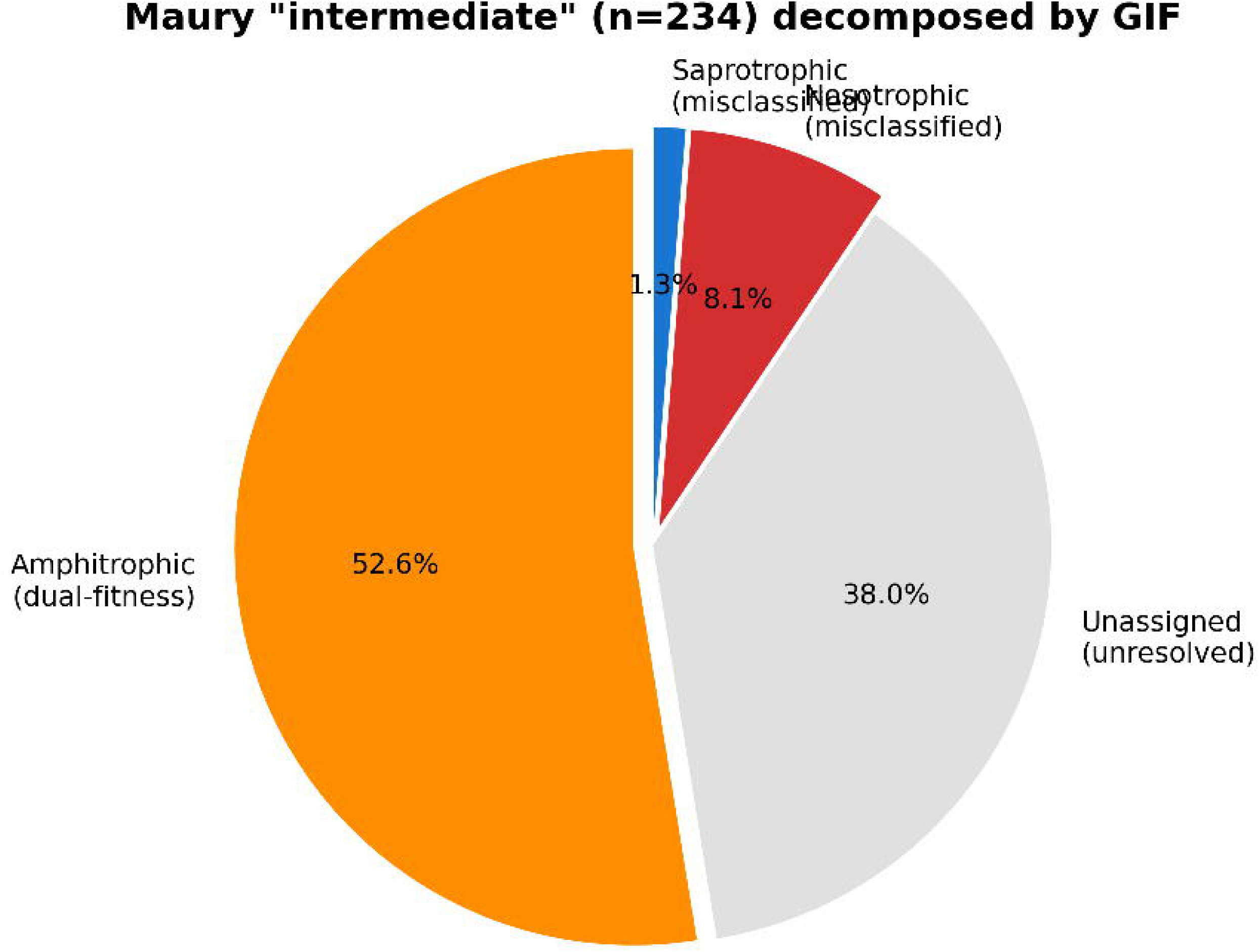

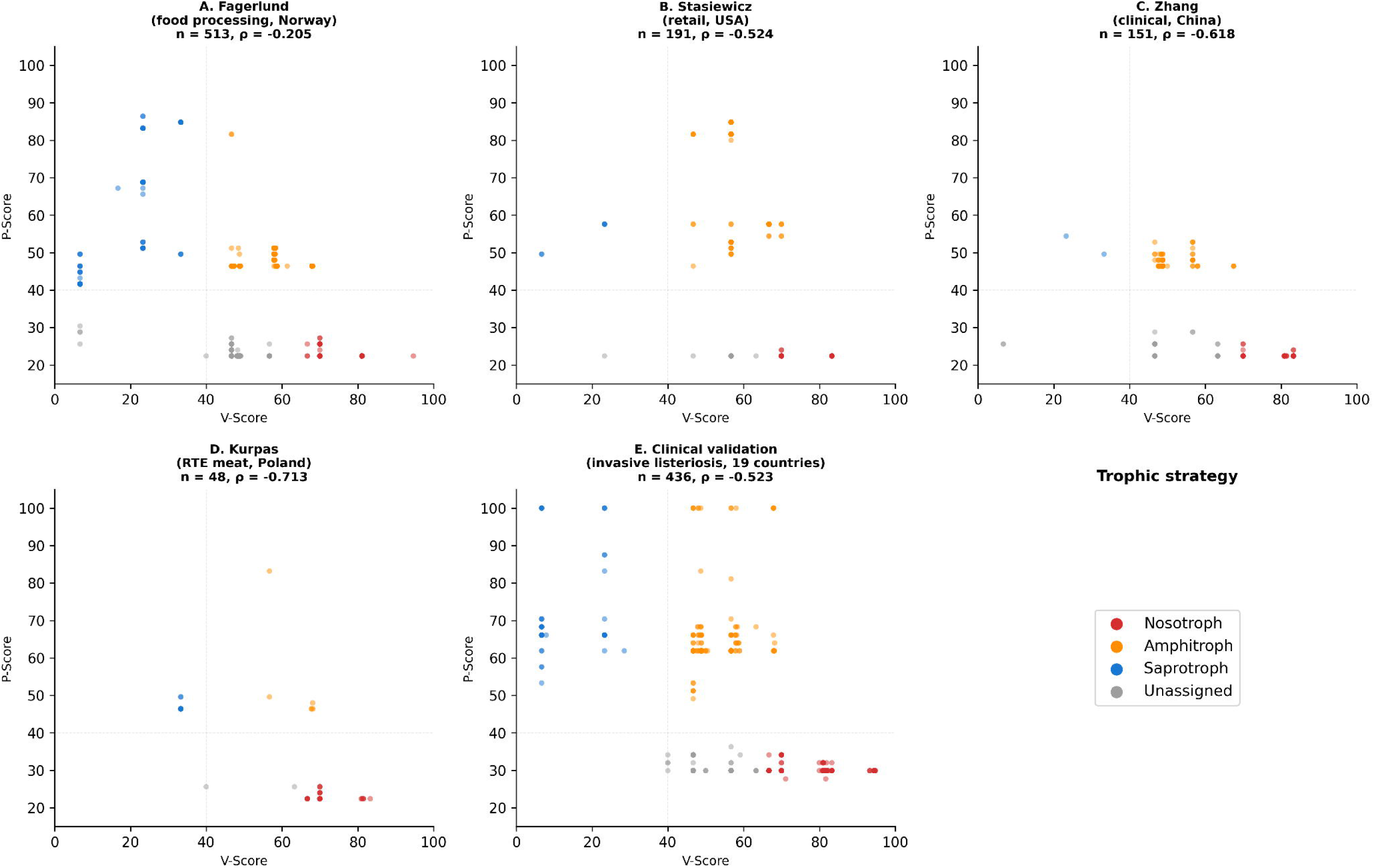

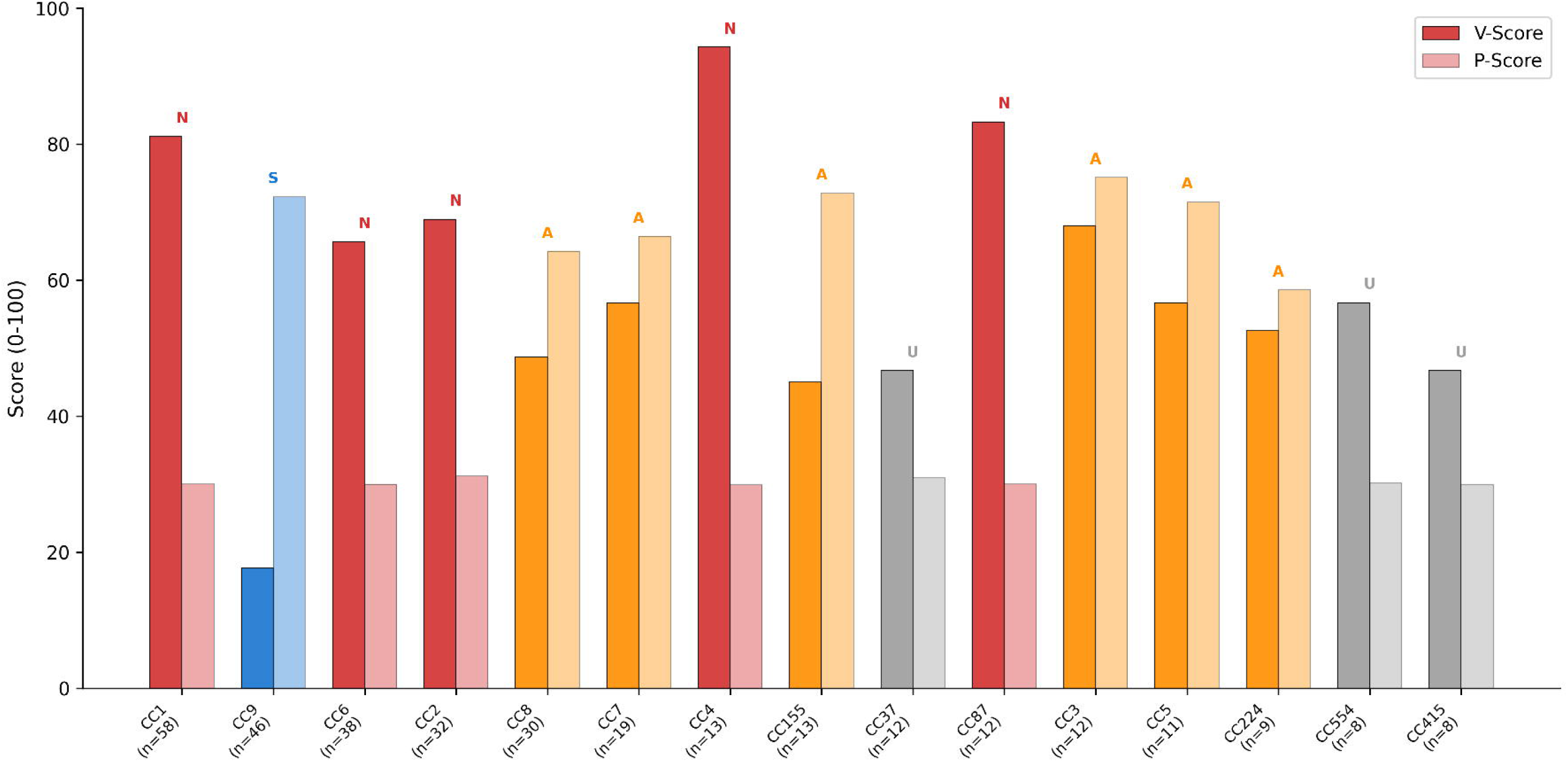

