## Supplementary data and tables for "Amphitrophic *Listeria monocytogenes* causes one-third of invasive listeriosis yet remains undetected by clonal complex-based risk classification"

### Supplementary Materials

*Amphitrophic Listeria monocytogenes constitutes one-third of invasive listeriosis: clinical validation of the Genomic Intelligence Framework exposes a systematic blind spot in genomic risk assessment*

Javier Gamboa — GIF Consortium, BIOTECNO, Spain

#### Supplementary Figure Legends

##### Figure S1. Virulence–persistence landscape across all GIF-characterised datasets

Scatter plots of V-Score versus P-Score for each dataset in the GIF validation corpus. Panels A-D show food-chain datasets from Gamboa (2026): (A) Fagerlund, n = 513, Norwegian food processing; (B) Stasiewicz, n = 191, US retail; (C) Zhang, n = 151, Beijing clinical; (D) Kurpas, n = 48, Polish RTE meat. Panel E shows the clinical dataset from this study (n = 436, invasive listeriosis). Points coloured by GIF trophic strategy: nosotrophic (red), amphitrophic (amber), saprotrophic (blue), unassigned (grey). Spearman rho indicated per panel. The aggregate negative V-P correlation at species level is replicated across all five datasets (rho = -0.205 to -0.713); within-strategy correlations are non-significant or positive; the only significant within-strategy signal is the amphitrophic positive correlation (rho = +0.221, p = 0.010).

##### Figure S2. GIF component scores by clonal complex

V-Score (solid bars) and P-Score (translucent bars) for the 15 most frequent clonal complexes in the clinical dataset. Bar colour indicates dominant trophic strategy: nosotrophic (red), amphitrophic (amber), saprotrophic (blue), unassigned (grey). Letters above bars indicate trophic classification (N/A/S/U). Sample sizes indicated below CC labels. Note that CC87 (n = 12, V-Score = 83.3) is classified as nosotrophic by GIF despite being "intermediate" under Maury et al. (2016), and CC9 (n = 46, V-Score = 17.7) is classified as saprotrophic despite causing confirmed invasive listeriosis.

#### Supplementary Tables

##### Table S1. Complete GIF results for 436 clinical isolates

Full GIF output (34 variables) for each of the 436 clinical L. monocytogenes isolates, including component scores (V, P, C, R), trophic strategy, virulence markers (InlA status, LIPI-1/3/4), persistence markers (qacH, bcrABC, SSI-1/2, cadmium), assembly quality metrics, and metadata (country, collection date, isolation source, SNP cluster, BioProject). Available as TSV file.

##### Table S2. CC-level statistics by trophic strategy

Summary statistics for each clonal complex (n >= 2) including mean and SD of V-Score and P-Score, dominant trophic classification, clinical and industrial GIF scores, and prevalence of key genomic markers (InlA functional, SSI-1, qacH, bcrABC, LIPI-3). Available as TSV file.

##### Table S3. Cross-tabulation of Maury epidemiological categories versus GIF trophic classification

Contingency table showing the distribution of GIF trophic strategies within each Maury epidemiological category (hypervirulent, intermediate, hypovirulent) for all 436 clinical isolates. Demonstrates that 47.4% of Maury "intermediate" isolates are not amphitrophic, comprising misclassified nosotrophs (8.1%), functionally unresolved lineages (38.0%), and misclassified saprotrophs (1.3%).

#### Supplementary Methods

##### S1. GIF scoring algorithm summary

The GIF score is computed as a weighted linear combination of four components: GIF = w_V * V + w_P * P + w_C * C + w_R * R, where weights differ by calibration context. Industrial calibration: w_V = 0.30, w_P = 0.40, w_C = 0.20, w_R = 0.10. Clinical calibration: w_V = 0.40, w_P = 0.20, w_C = 0.30, w_R = 0.10. Each component score ranges from 0 to 100. Full algorithmic specification is available at https://github.com/jgamboa-biotecno/GIF-Framework.

##### S2. Trophic strategy classification criteria

Trophic strategy is assigned per isolate based on V-Score and P-Score thresholds established and validated in Gamboa (2026): Nosotrophic: V-Score above virulence threshold AND P-Score below persistence threshold. Amphitrophic: both V-Score and P-Score above their respective thresholds. Saprotrophic: V-Score below virulence threshold AND P-Score above persistence threshold. Unassigned: neither criterion fully met. Thresholds are defined in the GIF specification and calibrated against the Fagerlund longitudinal dataset (AUC = 0.933).

##### S3. Bayesian imputation for P-Score Level 2

Clinical isolates lack temporal facility metadata required for P-Score Level 2 (operational persistence assessment). For these isolates, P-Score Level 2 is estimated via Bayesian imputation from genetic persistence markers (qacH, bcrABC, SSI-1, SSI-2, cadmium resistance operons, CRISPR absence). The imputation model was calibrated against the Fagerlund dataset where both genetic markers and temporal persistence data are available, achieving AUC = 0.980 for persistence classification. Imputed P-Scores are flagged in the output (Bayesian_imputation_used = YES) to maintain transparency.

##### S4. Dataset selection and stratification

Clinical isolates were retrieved from NCBI Pathogen Detection (March 2026) using the following filters: Organism = Listeria monocytogenes, Isolation type = clinical, Assembly = GCA_* (GenBank assembly accession), SNP cluster = assigned. From 20,219 qualifying records, stratified sampling was performed to ensure: (1) proportional representation across serogroups 4b, 1/2a, 1/2b, 1/2c and unknown; (2) geographic diversity with per-country cap of 30% and minimum 10 countries; (3) confirmed invasive source (blood, CSF, placenta) prioritised. Two assemblies failed NCBI download, yielding 436 for analysis. All 436 passed GIF quality control (100% completion rate). The final dataset spans 57 clonal complexes, 19 countries, 34 BioProjects, and collection dates from 1981 to 2025.
